# Spine-brain dynamics during human defensive threat-reactions

**DOI:** 10.1101/2025.07.18.665552

**Authors:** Elena Andres, Lycia D. de Voogd, David Norris, José Marques, Karin Roelofs

## Abstract

Synchrony between central and autonomic nervous system responses during threat rely on spine-brain interactions. Activation of the parasympathetic autonomic nervous system during acute threat has been linked to optimized action preparation and longer-term stress-resilience. However, it is unclear how this is orchestrated by spine-brain interactions during our widely used models of threat conditioning. Here we used a well-established threat conditioning procedure to test whether freezing-related bradycardia modulates the strength of motor- and sensory-relevant spinal-brain connectivity patterns. Conditioning elicited the expected autonomic and psychophysiological responses, with significant bradycardia and heightened differentiation between conditioned stimuli. Heart rate bradycardia modulated activity in the cingulate cortex and spinal cord, emphasizing their role in freezing. Notably, bradycardia-modulated spinal cord activity during threat was functionally coupled with motor cortex activity, suggesting a preparatory response for defensive actions. Establishing these effects for the first time in humans helps discovery of biomarkers for threat coping-relevant body-brain interactions, essential for clinical evaluations.

## Introduction

The ability to respond effectively to threat is a cornerstone of survival. Among the most critical adaptations is freezing, a state of immobility and bradycardia accompanied by heightened sensory awareness and motor readiness, which allows organisms to prepare for action while avoiding detection by predators ^1–8^. Although the relevance of threat-related freezing and bradycardia has been widely acknowledged for optimized threat coping ^5,6,9^, it remains unclear how spine-brain interactions facilitate action preparation under threat.

Here we use a well-established threat conditioning paradigm to test whether freezing-related bradycardia modulates the strength of motor- and sensory-relevant spinal-brain activity and connectivity. Testing spinal-brain threat-responses concurrently, during the most important and most widely used model of associative threat learning on a 3T MRI scanner, is relevant to provide a new widely applicable venue for future assessment of integrated body-brain biomarkers, that may be relevant for stress-related disorders.

Animal studies have indicated that freezing is governed by an interconnected defense circuit involving subcortical and cortical brain regions. Subcortical structures such as the amygdala and periaqueductal gray (PAG) are critical for initiating freezing, while cortical regions such as the anterior cingulate cortex (ACC) and motor cortex modulate freezing onset and offset during threat and safety learning ^10–15^. These neural defense circuitries have also been implicated in freezing-related bradycardia in humans ^3,14,16^.

The spinal cord plays a pivotal role in translating these neural signals into motor and autonomic outputs, integrating descending inputs from the brain with ascending sensory information in both animals ^12,13^ and humans ^17–21^. Although recent pioneering studies in humans have started to explore spinal-brain interactions during pain and emotion processing ^21–24^, no studies have investigated spine-brain connectivity during threat conditioning, let alone during freezing-related bradycardia. This represents a clear gap in understanding how central and peripheral systems dynamically coordinate during threat.

Freezing-related bradycardia refers to a unique psychophysiological state that is optimal for perceptual sensitivity and action preparation ^7,25,26^, for a review see Roelofs and Dayan (2022)^6^. Several studies have shown that individuals respond to threat of shock by concurrent activation of the sympathetic and parasympathetic branches of the autonomic nervous system ^5^. Sympathetic arousal increases action readiness and galvanic skin response, heart rate, and blood pressure. During freezing, concurrent -and even dominant-parasympathetic activation results in net bradycardia and a general break on the motor system ^6,27^. Because heart rate (HR) is predominantly parasympathetically innervated and skin conductance response (SCR) predominantly sympathetically, the defensive state of freezing is typically reflected in a net HR deceleration or bradycardia responses, with SCR largely preserved ^16,28–33^. Together, these patterns provide a distinct autonomic signature of the freezing state and reflect the underlying neural coordination of defensive behavior.

Recent human investigations have shown that freezing-related bradycardia is linked to enhanced sensory upregulation and precision ^3,29,33–35^ and action preparation ^5,27^. Several studies have shown that the stronger the magnitude of the HR deceleration under threat of shock, the faster one is for subsequent accurate responses ^3,29,31,32^. Such immediate physiological responses may reflect broader regulatory capacities relevant to long-term adaptation. Freezing is a reaction consistently observed during threat conditioning, a model widely used to study the processes underlying threat learning and anxiety ^2,36,37^. There is increasing evidence that the ability to exhibit freezing and parasympathetic dominance, reflected in HR bradycardia and heart rate variability (HRV), is linked to longer-term resilience ^30,38–42^ and that individuals with an anxiety-related disorder fail to show these patterns, presumably due to chronic sympathetic overactivation and reduced vagal tone ^43–46^. This makes it extra relevant to study the relation between freezing-related bradycardia and neural mechanisms involved in threat coping.

Although the relevance of threat-related freezing and bradycardia has been widely acknowledged for optimized threat coping ^5,6,9^, the interaction between autonomic and central systems under threat remains poorly understood. Specifically, it is unclear how freezing-related bradycardia influences spinal-brain connectivity and how this interaction contributes to sensory-motor integration and action preparation. Addressing these questions is critical for understanding the mechanisms underlying adaptive threat responses and their dysregulation in anxiety-related disorders.

We used a well-established threat conditioning paradigm combined with simultaneous brain and spinal cord imaging to investigate how freezing-related bradycardia modulates brain-spine neural activity and connectivity. By incorporating heart rate as a parametric modulator, we aimed to identify regions and pathways involved in freezing-related sensory-motor integration. Concretely we predict freezing-related bradycardia to be linked to increased motor spinal cord (ventral-SpC) – motor brain connections. In addition, a pattern of sensory spinal cord (dorsal-SpC) – sensory brain regions is expected.

## Results

### Threat conditioning successfully elicits threat-bradycardia

The main aim of the study was to determine whether freezing-related bradycardia modulates neural activity and connectivity across the spinal cord and brain, supporting its role as a central component of threat-induced sensory-motor integration.

Participants (n = 37; see Methods) underwent a threat conditioning (acquisition and extinction) paradigm while undergoing simultaneous brain and spinal cord fMRI. Two snake images served as conditioned stimuli (CS+ and CS−). The CS+ was paired with an aversive electric shock (reinforcement rate of 37.5%), while the CS− was never paired with an aversive electric shock (Fig. 1A). Simultaneous acquisition of brain and spinal cord activity was enabled by a specialized imaging sequence ^47^, covering spinal levels from approximately C3 to C6. Threat conditioning was successful as indicated by the typical patterns in sympathetic, subjective and neural activity patterns (see Fig. 1).

**Figure 1.**
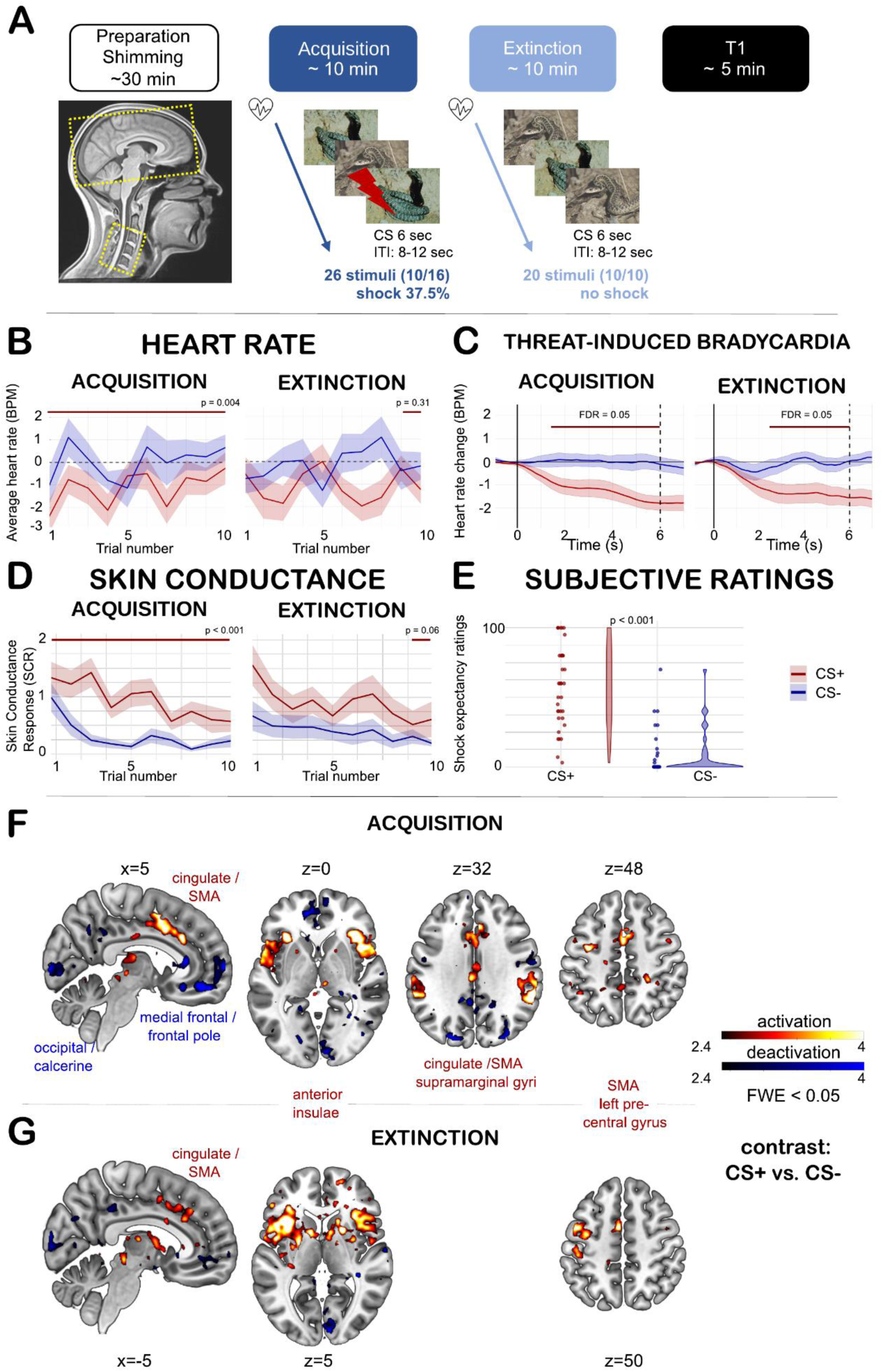
Experimental design and psychophysiological and neural features of threat conditioning. **A.** After preparation of the sequence ^47^, participants underwent a resting state and a threat of shock task (not shown in pipeline) followed by a threat conditioning paradigm. Lastly, an anatomical image was acquired. **B.** Trial-by-trial heart rate responses towards CS+ and CS-stimulus presentations in acquisition and extinction show successful conditioning. **C.** Trial-based heart rate across the 6-second trial period averaged for all trials reveal threat-induced bradycardia during acquisition towards the CS+ but not the CS-. The red line marks the time points where differences were statistically significant (p<0.05, FDR-corrected). Vertical lines indicate stimulus onset (0 seconds) and offset (6 seconds). **D.** As far as sympathetic responses, trial-by-trial skin conductance responses (SCR), were significantly greater for CS+ than CS− during both acquisition (W = 95, p < .001) and extinction (W = 98, p < .001, Fig. 1D).

Critical for here was to confirm successful freezing-related bradycardia patterns during conditioning, particularly focusing on parasympathetic dominance reflected in HR deceleration. Heart rate responses were quantified on a trial-by-trial level as the average baseline-corrected HR over a 6 s window after stimulus onset, relative to a 1 s pre-stimulus baseline (see Methods), with lower values indicating stronger bradycardia. We found significantly lower mean heart rate (BPM) for CS+ compared to CS-during acquisition (W = 421, p = .004, Fig. 1B trial-by-trial, C: trial-based) and extinction (W = 419, p = .004), indicative of a freezing-like bradycardia state. The last two extinction trials revealed no significant differences between CS+ and CS-responses (W = 590, p = .311, Fig. 1C), suggesting successful extinction. These results confirm that the conditioning paradigm effectively elicited a freezing-like bradycardia response during acquisition that diminished with extinction learning. This effect was further confirmed by the fact that net bradycardia was accompanied by concurrent sympathetic responses (see SCR in Figure 1D and legend Figure 1).

### Freezing-related bradycardia modulates neural activity in spinal cord and brain

Having confirmed that threat conditioning was successful and produced a robust freezing-related bradycardia response, we next investigated whether this parasympathetic state was associated with specific neural activation patterns in the spinal cord and brain.

To investigate spinal correlates of freezing, we conducted a time-sensitive Finite Impulse Response (FIR) analysis (Fig. 2A). This approach modeled HR-modulated activity of CS+ versus CS- in time bins corresponding to the repetition time (TR = 3.02 s), starting one TR before stimulus onset (see FIR without HR-modulation in Extended Data Fig. 1). During acquisition, non-reinforced CS+ trials elicited greater HR-modulated activity at stimulus offset compared to CS− in the spinal cord, with effects observed in the whole grey matter (estimated effect = 0.01, 90% CI [0.00, 0.02], posterior probability = 0.97, evidence ratio = 35.81), as well as separately in the dorsal (estimated effect = 0.01, 90% CI [0.00, 0.02], posterior probability = 0.94, evidence ratio = 15.85) and ventral regions (estimated effect = 0.01, 90% CI [0.00, 0.02], posterior probability = 0.95, evidence ratio = 17.29). These patterns are consistent with enhanced sensory precision and motor readinesss associated with freezing. Importantly, these effects were specific to acquisition. No significant HR-modulated differences between CS+ and CS− were observed in the spinal cord during extinction, highlighting the task-specific nature of these responses.

**Figure 2.**
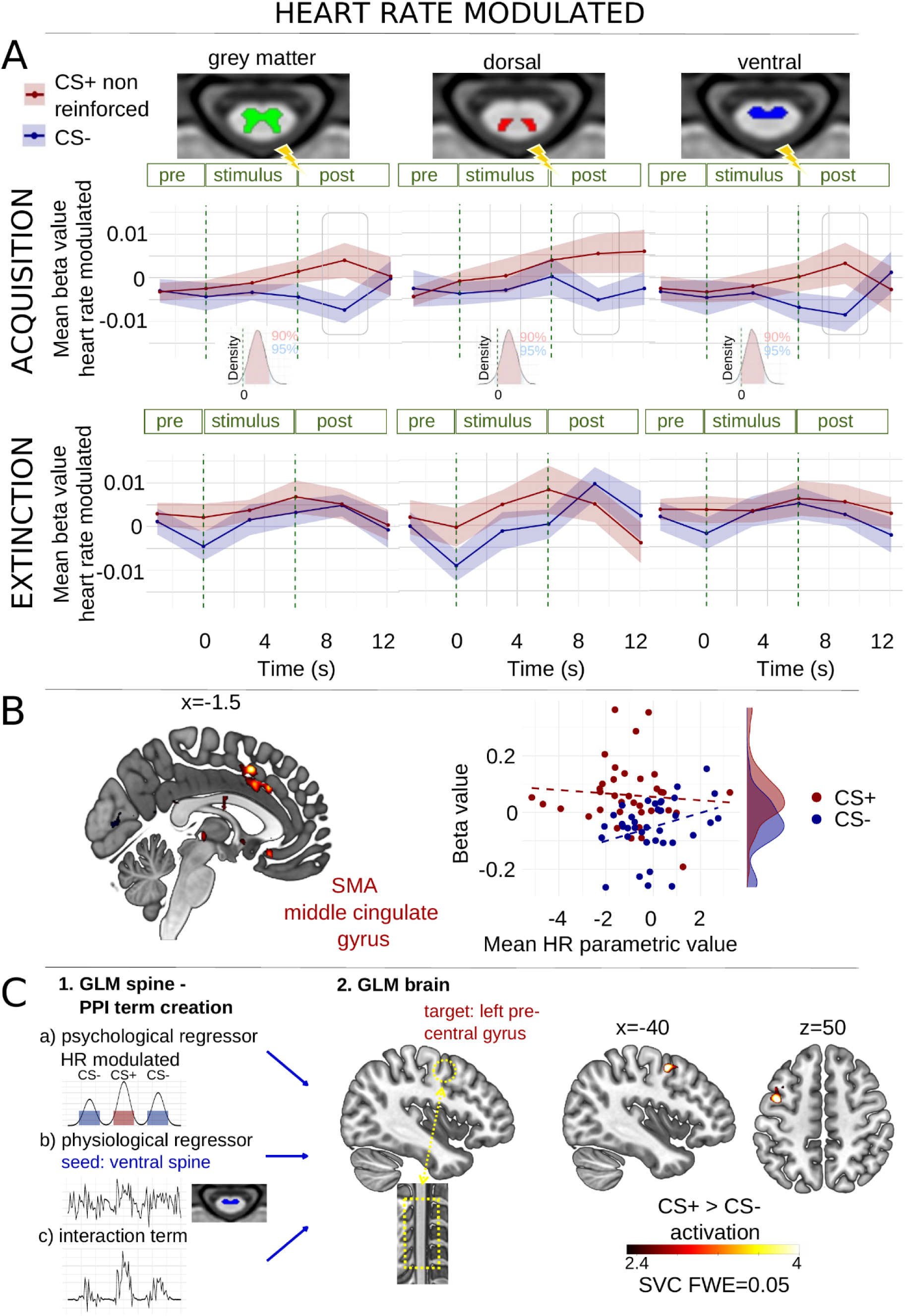
Heart-rate modulated activity towards CS+ and CS- and spine-brain coupling. **A**. Heart-rate modulated spinal cord grey matter (left), dorsal (mid) and ventral (right) responses starting one TR before stimulus onset revealed a significant increase in CS+ responses at the TR of stimulus offset compared to CS-responses during acquisition (grey boxes), whereas this effect was abolished in extinction. **B.** Heart-rate modulated significant cluster in SMA/middle cingulate gyrus and beta estimates separately for CS+ and CS-parametric regressors in relation to their averaged parametric value. A negative HR parametric value represents bradycardia. Lines represent linear regression lines for each stimulus type. **C.** A PPI taking the ventral spinal cord as a seed during stimulus presentation revealed coupling to the left precentral gyrus. SVC = small volume correction applied.

To assess brain responses specifically associated with freezing-related bradycardia, we included trial-by-trial HR values as a parametric modulator in the GLM for CS+ and CS−trials. Significant differences in HR-modulated brain activity for our stimulus types were observed during acquisition but not extinction (Fig. 2B) in the SMA / cingulate cortex (CS+ > CS-: estimated effect = 0.106, SE = 0.027, t(70) = 3.996, p < .001). The main effect of the parametric HR values (estimated effect = 0.025, SE = 0.013, t(70) = 1.838, p = .070) and interaction between HR and CS type (estimated effect = −0.031, SE = 0.017, t(70) = −1.785, p = .079), were not significant (but showed a trend in the expected direction).

Together, these results demonstrate that the physiological state of freezing was reflected in the neural dynamics of both the spinal cord and the brain, with significant responses observed during acquisition but not extinction. The findings suggest a modulation of SMA / cingulate cortex activity during threat (vs safety) processing when trial-by-trial fluctuations in freezing-related bradycardia are taken into account.

### Freezing-related bradycardia is associated with spinal–cortical motor connectivity

To test our main hypothesis, we next examined how bradycardia-related activity in the spinal cord is functionally integrated with brain activity during threat anticipation. We conducted a psychophysiological interaction (PPI) analysis using HR-modulated spinal cord activity as the physiological regressor.

A GLM of the spinal cord data during acquisition was created, incorporating CS+ and CS− conditions as well as trial-wise HR parametric modulation (see control PPI of GLM with CS offsets corresponding to significant FIR timepoints in Extended Data Fig. 2). From this model, we extracted HR-modulated activity from grey matter, dorsal, and ventral spinal cord volumes of interest (VOIs), adjusted for nuisance regressors. These were used to construct PPI terms, which were then entered into a second-level GLM for brain activity.

Three a priori regions of interest (ROIs) were targeted: the cingulate cortex and the motor cortex (as identified in the brain GLM without HR parametric modulation), and the periaqueductal gray (PAG), using small-volume corrections (SVC).

Significant HR-modulated activity was observed in the left motor cortex (Fig. 2C), which was functionally coupled with HR-modulated activity in the grey matter and ventral spinal cord during CS+ compared to CS− (grey matter SVC: xyzMNI: -40/0/50, T = 4.75, kE=32, p_FWEcorr_=.005, whole-brain: kE=423, p_FWEcorr_=.003; ventral SVC: xyzMNI: -44/3/51, kE=32, p_FWEcorr_=.005; whole-brain: kE=258, p_FWEcorr_=.058).

These results suggest that during threat anticipation, bradycardia-related activity in the spinal cord — particularly in ventral spine associated with motor function — was differentially coupled with left motor cortex activity.

However, during the last two extinction trials, this difference was no longer significant (W = 199, p =.062), suggesting successful extinction learning. **E.** Subjective ratings of shock expectancy collected at the end of both conditioning tasks revealed significantly higher ratings for CS+ compared to CS− (W = 77, p < .001; Fig. 1E), confirming explicit awareness of the threat-safety association and successful acquisition of the conditioned contingency. **F.** A whole-brain general linear model (GLM) confirmed typical neural activation patterns associated with conditioned threat (Fig. 1F,G; Table 1). During acquisition, greater activation for CS+ than CS− was observed in the cingulate cortex, supplementary motor area (SMA), bilateral anterior insulae, and supramarginal gyrus. Subthreshold activations were observed in the left (p_uncorr_ = .018, xyzMNI = -34/-6/48, T = 5.16, kE = 102) and right precentral cortex (p_uncorr_ = .055, xyzMNI = 44/0/56, T = 4.36, kE = 63), with the left activation aligning with the motor region contralateral to the hand that received stimulation. Reduced activation for CS+ relative to CS− was found in the occipital/calcarine cortex and medial frontal cortex/frontal pole. **G.** During extinction, similar regions (cingulate/SMA, anterior insulae, and left precentral cortex) were again activated, consistent with their role in threat and safety processing (Fig. 1G, see Table 1 for whole-brain cluster level corrected inferential statistics). All labelled areas are significant at p < .05 FWE-corrected.

**Table 1.** Peak voxel coordinates and cluster statistics. All coordinates are defined in MNI152 space. All reported statistics are significant at p < .05, cluster corrected.

| | Region | Side | x (mm) | y (mm) | z (mm) | $p_{\text{corr}}$ | T | $k_E$ |
| --- | --- | --- | --- | --- | --- | --- | --- | --- |
| **Acquisition**<br>**CS+ > CS-** | Supramarginal gyrus | R | 60 | -40 | 34 | <.0001 | 7.04 | 1818 |
|  |  | L | -62 | -27 | 27 | <.0001 | 4.68 | 616 |
|  | Anterior insula | R | 42 | 16 | 4 | <.0001 | 6.67 | 2649 |
|  |  | L | -57 | 8 | 3 | <.0001 | 5.47 | 1029 |
|  | Anterior / middle cingulate gyrus | R/L | 3 | 12 | 38 | <.0001 | 5.37 | 1029 |
| **CS+ < CS-** | Medial frontal cortex | R / L | -12 | 57 | 24 | <.0001 | 5.62 | 1950 |
|  | Middle / inferior occipital gyrus | R | 45 | -74 | 2 | <.0001 | 5.51 | 1134 |
|  | Hippocampus | R | 18 | -12 | -26 | .028 | 4.69 | 300 |
|  | Calcerine cortex | R | 18 | -88 | -4 | <.0001 | 4.52 | 571 |
|  | Posterior cingulate gyrus | R / L | -6 | -50 | 38 | .019 | 4.31 | 322 |
| **Acquisition HR modulated CS+ > CS-** | Middle cingulate gyrus / supplementary motor area | R / L | -2 | 12 | 46 | .022 | 4.17 | 291 |
| **CS+ < CS-** | Calcerine cortex | R | 9 | -84 | 4 | .012 | 5.20 | 323 |
| **Extinction CS+ > CS-** | Anterior insula | L | -39 | 16 | 3 | <.0001 | 6.20 | 3488 |
|  |  | R | 36 | 22 | 0 | <.0001 | 5.46 | 2292 |
|  | Middle cingulate gyrus / supplementary motor area | R / L | -4 | 0 | 48 | .001 | 5.69 | 493 |
|  | Dorsal striatum | R | 22 | -6 | 9 | <.0001 | 5.68 | 648 |
|  | Precentral gyrus | L | -40 | -6 | 54 | .002 | 5.20 | 465 |
| **CS+ < CS-** | Middle occipital gyrus | R | 38 | -75 | 38 | .002 | 4.76 | 445 |
|  | Calcerine cortex | R | 8 | -92 | 3 | .003 | 4.37 | 429 |
|  | Anterior cingulate gyrus | R / L | 0 | 46 | -4 | .007 | 4.23 | 375 |

## Discussion

### Overview

This study implements simultaneous brain–spine imaging during threat conditioning to explore the neural basis of body–brain interactions in defensive reactions. The use of a well-established threat conditioning model allows us to extend a wealth of current knowledge by demonstrating that freezing-related bradycardia is not only reflected in autonomic changes but also modulates activity within the spinal cord, in a way that it is synchronous to neural changes. Concretely, we observe that threat anticipatory bradycardia during acquisition is associated with 1) increased activity in the ventral and dorsal regions of the spinal cord and 2) neural motor regions, -and most critically-3) is associated with increased connectivity between the ventral spinal cord and motor cortex. These findings point to an integrated mechanism of action preparation during threat, suggesting that even in the absence of overt behavior, the body engages in preparatory processes to optimize defensive responses.

### Neural signatures of freezing

Freezing-related bradycardia was accompanied by distinct neural signatures. In the spinal cord, activity at stimulus offset increased as a function of trial-by-trial bradycardia in dorsal and ventral regions, suggesting coordinated recruitment of motor and sensory pathways during threat anticipation. In the brain, freezing-related bradycardia was associated with activity in the SMA / cingulate cortex, a region implicated in the control and maintenance of defensive states. Crucially, bradycardia-related activity in the ventral spinal cord was functionally coupled with the left motor cortex, consistent with our prediction of integrated motor processing across spinal and cortical levels. Together, these findings support the idea that parasympathetic freezing is implemented through distributed neural dynamics that span brain–spine circuits involved in sensory–motor integration ^6,11^.

### Extending freezing models to the spine

Building on research characterizing freezing as a parasympathetically dominated defensive state, our results add an important dimension by directly linking autonomic changes to spinal cord activity. Previous studies ^3,14,16^ have described freezing as attentive immobility involving sensori-motor and cingulate–brainstem pathways that prepare the organism for optimized action ^6^. Freezing-related bradycardia and cingulate/motor activity have been linked to fast and value-based motor responses under threat, respectively ^31^.

Freezing-related bradycardia has also been linked to perceptual sensitivity ^29,33,34^, suggesting that the physiological state of freezing involves a shift in sensory processing toward rapid threat detection. Our data extend these processes to the spinal level, where increased activity at stimulus offset may indicate fine-tuning of sensory inputs for action preparation.

### Spinal cord specificity and extinction effects

Crucially, this effect was absent during extinction, indicating that spinal cord responses are not merely a byproduct of stimulus presentation but are shaped by threat contingencies. The disappearance of this effect during extinction suggests that the spinal cord plays an active role in encoding threat associations rather than passively reflecting autonomic changes. These findings highlight the spinal cord as an integral component of the threat response network, dynamically engaged in the regulation of defensive states based on learned threat predictability.

### Freezing, decision-making, and autonomic modulation

The integration of heart rate modulation with spinal cord responses also provides a potential mechanism by which freeze-states may impact decision making under threat. Klaassen et al. ^31,32^ showed that freezing-related bradycardia affects approach–avoidance decisions by modulating the impact of subjective value on choice behavior and may mediate approach–avoidance arbitration. Likewise, Hashemi et al. ^3^ demonstrated that midbrain activity, including the PAG, facilitates the shift from freezing to rapid action execution. These studies highlight that autonomic states not only prime sensory–motor systems but also shape higher-level decision processes. Our finding of heart rate– modulated brain–spinal coupling underscores the possibility that autonomic signals serve as a bridge between the evaluation of threat and the preparation of defensive action.

### Clinical future direction for anxiety-related disorders

These findings provide new starting points for the identification of neural defense circuits alteration which may underly maladaptive threat-coping in anxiety disorders. Previous investigations into the effects of aversive life events on freezing responses ^44^ and the predictive value of early freezing behavior for later internalizing symptoms ^39^ highlight the clinical relevance of understanding these mechanisms. By demonstrating that freezing-related autonomic changes are coupled with specific patterns of spinal cord activity, our study provides a potential biomarker for the integrity of body–brain integration during threat, which may be disrupted in anxiety and stress-related disorders.

### Expanding spinal cord imaging to anticipatory threat and action preparation

Previous studies using combined brain–spinal cord imaging in humans have primarily focused on sensory modulation during pain and emotional processing . For example, work on nocebo hyperalgesia ^48^, attentional analgesia ^23^, and social observation of pain ^24^ has revealed that cognitive and affective states modulate spinal activity. Similarly, spinal-only fMRI studies have shown that emotional arousal ^17,19^ and aversive sensory input ^18^ modulate spinal activity, supporting the view that affective and motivational states shape spinal processing independently of direct pain stimulation. Additionally, work by Vahdat et al. (2015) ^49^ demonstrated that spinal–cortical connectivity is shaped by motor learning, highlighting the functional plasticity of the spinal cord in adaptive behavior. Our study extends this work by demonstrating, for the first time, that an autonomic state, i.e. freezing-related bradycardia, modulates spinal–cortical connectivity during threat learning. While pain and threat engage distinct processes, they both serve as potent signals of potential harm and activate overlapping defensive systems. Our results suggest, that spinal–cortical dynamics are not limited to processing actual aversive input, but also play a role in anticipatory responses to potential threat. We thereby broaden the scope of spinal imaging research to include learned threat states that precede physical harm.

### Interpretational issues

While we found robust coupling between heart rate modulation and spinal cord-motor connectivity, we did not observe significant connectivity with other regions typically associated with freezing, such as the PAG. On the one hand this confirms the important role of (primary) motor cortex in freezing ^11^. On the other hand, future studies should consider employing higher temporal resolution to increase sensitivity in small midbrain regions ^50–52^. Indeed, recent research has sparked renewed interest in the role of the primary motor cortex in freezing-related processes. For instance, Bai et al. (2023) ^11^ demonstrated its direct involvement in the modulation of defensive states, suggesting a more active role than previously assumed. Importantly, anatomical studies have shown that the primary motor cortex maintains direct corticospinal projections, allowing for a fast and efficient relay of motor commands to the spinal cord ^53–56^. In contrast, regions like the PAG and the cingulate cortex, influence spinal processing more indirectly—likely via relays through the brainstem or medulla ^57,58^. This difference in structural connectivity may explain the robust HR–modulated coupling observed between the spinal cord and motor cortex in our data, and the absence of significant connectivity with midbrain regions such as the PAG. It also reinforces the view that the primary motor cortex plays a pivotal role in the preparatory aspects of freezing by engaging directly with spinal circuits to prime the body for action.

A methodological aspect of our imaging protocol is the relatively long repetition time (TR) of 3.02 seconds. In our design, each TR sequentially captures brain activity for approximately 2 seconds, followed by spinal cord activity for about 1 second. Because stimulus onset was not locked to the beginning of a TR and stimulus length was 6 sec, stimulus offset typically fell within the third TR. This aligns with the observed heart rate– modulated activity in the grey matter, ventral, and dorsal spinal cord regions. Notably, the pronounced activity at the third TR is consistent with the anticipated temporal profile of the task, coinciding with the period when a shock is expected and preparatory processes are engaged.

## Conclusion

In sum, our findings provide new evidence that the autonomic changes during threat conditioning are closely linked to spinal cord activity and its connectivity with brain regions involved in motor planning and sensory processing. Proven modulations by threat and autonomic states linked to robust activity in the ventral as well as dorsal spinal regions were specific for threat acquisition and disappeared with extinction, stressing the specificity of the findings. Establishing these effects for the first time in humans is relevant to provide a biomarker for threat coping-relevant body-brain interactions. In addition, by establishing spinal-brain threat-responses on a 3T MRI scanner opens the way for wide application of relevant body-brain interactions during threat conditioning. These results not only deepen our understanding of the neural substrates underlying defensive reactions but also offer a potential biomarker for the integrity of body–brain interactions that could inform future research into anxiety and stress-related disorders.

## Materials and methods

This study was preregistered on AsPredicted (https://aspredicted.org/3V1_7SC). All research activities were carried out in accordance with the Declaration of Helsinki and approved by the local ethics committee (Ethical Reviewing Board CMO/METC [Institutional Research Review Board] Arnhem-Nijmegen, CMO 2014/288) and conducted according to these guidelines and regulations (i.e., medical/scientific research).

### Participants

Fifty-five healthy adult participants were recruited for this study based on the following inclusion criteria: fluency in English; no history of psychiatric or neurological disorders; no regular medication use except contraceptives; and no MRI contraindications.

Seven participants did not complete the study for the following reasons: early termination by the participants (n=3); poor signal quality during acquisition (n=3), and technical issues (n=1). Furthermore, nine participants were excluded before analysis due to poor data quality based on MRIQC ^59^ output inspection (brain, n=7), and visual inspection of spine data (n=5, out of which 3 were also excluded during brain data inspection). Lastly, two participants were excluded based on insufficient HR data during both tasks, and one more participant was excluded only in the extinction analysis based on HR data quality.

Thus, the final sample consisted of 37 participants for acquisition and 36 for extinction analyses (15 female, 22 male, mean age ± standard deviation [SD] = 27 ± 10.6). For more detailed demographic information, see Table 1. Participants signed an informed consent before participating and received monetary reimbursement (26.25 Euro) for their participation.

**Table 1.** Demographic information (n = 37)

|  |  |
| --- | --- |
| Sex | 15 females, 22 males |
| Age (mean ± SD) | $27 \pm 10.5$ |
| Ethnicity | 32 White, 4 Asian, 1 Black or African American |
| BDI-II score (mean ± SD) | $1.8 \pm 2.4$ (min – max = 0 – 21) |
| STAI-T score (mean ± SD) | $45.8 \pm 4.7$ (min – max = 20 – 80) |
| CTQ score (mean ± SD) | $35.8 \pm 10.0$ (min – max = 25 – 125) |

### Study Design

The total duration of the overall study was approximately 100 minutes (Fig. 1). Upon arrival, participants completed MRI screening and filled out questionnaires containing demographic and mental health inquiries.

Subsequently, participants were positioned in the MR scanner and prepared for physiological monitoring during scanning (respiration belt, pulse oximeter, skin conductance electrodes, see section **Peripheral stimulation and measurement**). Electrodes for electric shock delivery were attached. During the MR shimming procedure for the corticospinal imaging protocol (30 to 45 min) ^47^, participants were instructed to relax and had the option to view a nature documentary. This was followed by a standardized 5-step shock work-up to calibrate the intensity of the electric shock which remained the same for the rest of the task (see section **Peripheral stimulation and measurements**).

Next, a resting state, threat of shock task, followed by the threat acquisition and extinction tasks were performed. Data presented in this manuscript only refers to the tasks of acquisition and extinction.

Participants were given verbal instructions which were also available on the screen before acquisition, indicating two consecutive rounds of the task. After the experimenter verified that instructions were understood, the experimental paradigm was started. Acquisition lasted approximately 8 minutes (145 volumes), while extinction took around 5 minutes (114 volumes) to complete. Finally, a T1-weighted anatomical scan of the brain and the cervical spinal cord was acquired.

### Threat acquisition and extinction

During acquisition, pictures of two distinct snakes were employed as conditioned stimuli (CS) sourced from the International Affective Picture System ^60^. One snake image (CS+) was paired with an electric shock which served as an unconditioned stimulus (US) with a reinforcement rate of 37.5 %, while the other was never accompanied by a shock (CS-). Conditioning included 16 CS+ (six reinforced) and 10 CS-trials. Each stimulus was displayed for 6 seconds, with a jittered inter-trial interval (ITI) ranging from 8 to 12 seconds. During reinforced CS+ trials, the shock was administered 200 ms before the stimulus offset and co-terminated with stimulus offset. The first two stimuli served for image familiarization and were never paired with a shock, while the third stimulus was always accompanied by the US. During extinction, the same two images were presented each ten times without reinforcement. Images were assigned randomly and counterbalanced across participants.

Participants received verbal and written instruction before acquisition. They were instructed that one image might be followed by a shock while the other was always safe, but they were unaware of the specific associations. Participants were further told that the task would be executed in two runs. After acquisition, participants were informed that the next run would be following shortly. Following both experimental tasks, participants rated their expectancy of receiving a shock for each stimulus type on a continuous scale ranging from 0 to 100.

Acquisition and extinction tasks were executed with Expyriment, a Python library (version 3.7.16; ^61^), using script adjusted from Linda de Voogd (https://github.com/lindvoo/fcwml_mri_expriment).

### Peripheral stimulation and measurement

Physiological responses, including skin conductance response (SCR), heart rate (HR), and respiration, were recorded during MRI acquisition using a BrainAMP EXG MR 16 channel amplifier (resolution: 152.5 μV; sampling rate: 1000 hertz [Hz]) and BrainVision Recorder software (version 1.25.0001; Brain Products GmbH, 2020).

Two 10 mm Ag/AgCl electrodes connected to galvanic skin resistance sensors and coated with NaCl gel were attached to the participant’s left palm to record SCR. For HR measurement, a pulse wave sensor was placed at the participant’s left index finger, along with a hand glove to maintain warmth and optimize signal quality. Respiration was recorded via a respiration belt tied around the diaphragm of the participant.

Electric shocks were delivered via two electrodes attached to the fingertips of the index and middle fingers on the right hand (two 10 mm Ag/AgCl electrodes coated with gel for better adhesion, connected to a 9V MAXTENS 2000 shocker machine). For electric shock calibration, participants verbally rated the discomfort level of five consecutive shocks on a scale ranging from 1 (not uncomfortable at all) to 5 (very uncomfortable) after each shock. Shock intensity was adjusted based on these ratings so that it was uncomfortable but not painful. The first shock administered was always 8 volts (V) / 16 milliamperes (mA) and the intensity could vary from 0 to 40 V / 0 – 80 mA in 10 intensity steps (mean ± SD average intensity step: 3.9 ± 1.8; de Voogd et al., 2016).

### Questionnaires

Before the MRI session, participants filled out several questionnaires. First, they responded to a few demographic questions (general information, e.g. year of birth, sex, handedness). Subsequently, participants completed the short version of the Beck Depression Inventory II (BDI-II), the State-Trait Anxiety Inventory - Trait (STAI-T), and the Childhood Trauma Questionnaire (CTQ) ^62–64^. Notably, one question from the CTQ was missing in participant one to six due to a technical issue, which was not adjusted for in the demographic calculations (Table X) for this study. BDI values of 0–3 indicate minimal depression (n=31); 4–6 indicate mild depression (n=4); 7–9 indicate moderate depression (n=1); and 10–21 indicate severe depression (n=1). STAI-T values of 20-37 indicate no or low anxiety (n=2), values of 38-44 indicate moderate anxiety (n=14), and values of 45-80 high anxiety (n=21). Lastly, a cutoff value ≥ 35 for total CTQ scores indicates a significant history of childhood trauma (n=18).

### Physiological Data analysis

HR data were processed using in-house software for interactive visual artifact correction and peak detection, resulting in time-series of interbeat intervals expressed in beats per minute (BPM). Trial-by-trial quantifications of our psychophysiological index of freezing (i.e., bradycardia) were computed by extracting per trial the mean (HR) signal over a time window of 6 s and baseline corrected using a −1 s pre-stimulus onset average. These trial-by-trial indices were included as parametric modulators in the fMRI data analyses. To examine differences in heart rate responses between overall CS+ and CS-conditions over an 8-second trial period (6-second stimulus ± 1second), we performed a permutation test on the smoothed data. For each of 1100 time points, we computed the observed difference in mean heart rate between conditions across subjects. To assess the significance of these differences, we generated a null distribution by randomly permuting condition labels 1000 times and recalculating the mean difference for each permutation. This allowed us to estimate two-tailed p-values by comparing the observed differences to the null distribution. To control for multiple comparisons, we applied a False Discovery Rate (FDR) correction with an α level of 0.05.

After downsampling to 250 Hz, skin conductance responses (SCRs) were automatically scored with additional manual supervision using Autonomate ^65^ implemented in MATLAB (2023b) was employed, supplemented by additional manual supervision, to score response amplitudes of the SCR. The SCR response could start rising anywhere between 0.5 s and 6.5 s after stimulus onset, with a minimum rise time of 0.5 s and maximum rise time of 6 seconds, resulting in an overall response window of 12.5 seconds. In cases where multiple responses fulfilled these criteria, the largest response amplitude was taken into account. Responses smaller than 0.02 microsiemens (μS) were scored as zero and remained in our dataset. All trials reinforced with electric shock (CS-US) were omitted from analyses.

Before statistical analysis, additional participants were excluded from our sample based on the following criteria: no response to the US (electric shock) above 0.05 μS during conditioning (n = 3); mean response towards CS+ is smaller than towards CS-(n=5/9 at conditioning/extinction) or due to poor data quality (all zero responses, n = 7/12 at conditioning/extinction). This was done to be consistent with exclusion criteria previously used in our laboratory ^29^. Therefore, the final sample considered for statistical analysis of SCR consisted of 23/17 participants, in acquisition and extinction, respectively. All amplitude differences were square root transformed prior to statistical analysis.

### MRI data acquisition

We used a 3T MAGNETOM Prisma MRI scanner (Siemens AG, Healthcare Sector, Erlangen, Germany) together with a Siemens 64-channel head-neck coil to for image acquisition.

Head and neck movement were minimized by placing cushions on either side of the head. Additionally, adhesive tape was secured to the sides of the coil and participants’ foreheads to further restrict head motion and make patients aware of their initial position. The isocenter was positioned approximately at the participants’ chin, corresponding to the second and third cervical segments (C2-C3).

A specialized echo-planar imaging (EPI) sequence developed by Finsterbusch et al. (2013) was adapted to allow simultaneous acquisition of brain and spinal cord imaging data using two fields of view (FOVs) with distinct geometric resolutions and timing parameters. This setup allowed the optimization of imaging settings for each subvolume, improving overall image quality for both regions. To achieve simultaneous imaging of subvolumes with different geometries and timings, optimal shim parameters and resonance frequencies were determined for each subvolume before functional scanning. These parameters were dynamically adjusted during the recording of BOLD responses in the brain and spinal cord, respectively. Detailed information on the MRI pulse sequence and acquisition strategy can be found in Finsterbusch et al. (2013).

In the preparation phase, we first positioned the two slice groups as depicted in Fig. 1. The brain slice group consisted of 35 slices with a thickness of 3 mm and an in-plane resolution of 2x2 mm (FOV = 128 x 128 mm; flip angle = 75°). These slices were acquired in descending order along the anterior commissure - posterior commissure line (AC-PC line), covering most of the brain. The lower slice group for the spinal cord contained 9 slices perpendicular to the spinal cord, covering segments C3 - C6 (in-plane resolution = 1x1 mm; slice thickness = 5 mm + 1 mm gap; FOV = 128 x 128 mm; flip angle = 75°). Both FOVs utilized GRAPPA acceleration (factor 2) and flow re-phasing in the slice direction, resulting in echo times of 30 and 31 ms for brain and spinal cord slices, respectively.

Two saturation regions in a "V"-shaped orientation were applied to both sides of the FOVs to reduce ghosting and noise caused by blood flow in cervical vessels and breathing.

Before the scanning of functional images, we performed a shimming procedure to calculate optimal parameters for the brain and spinal cord FOVs. EPI volumes were acquired during this procedure to determine the best linear shim and resonance frequency values for each FOV separately, as well as for one combined FOV encompassing both the brain and spinal cord.

Additionally, a z-shimming procedure for the spinal cord slices was conducted to ensure low through slice dephasing in the presence of the large magnetic field inhomegeinities unduced by the irregular (and highly diamagnetic) cervical bone structure. This involved acquiring spinal cord images with 21 equidistant gradient steps, capturing slices with slight variations in the gradient strength or direction. Among these images, we identified the gradient setting resulting in the highest signal intensity within the spinal cord for each slice.

Acquisition times for the brain and spinal cord subvolumes were 2320 and 700 ms, resulting in an overall repetition time of 3020 ms for all slices. DICOM files were converted into the NIfTI format resulting in separate datasets for brain and spinal subvolumes.

High-resolution T1-weighted anatomical images were acquired using an MPRAGE sequence (TR = 2300 ms; TI=1100ms, TE = 3.03 ms; in-plane resolution = 1x1 mm; slice thickness = 1 mm; FOV = 256 x 225 x 192; flip angle = 8°), covering head and neck up to the upper thoracic vertebrae.

### Preprocessing fMRI data

Brain and spinal data were processed separately to address the specific requirements of each subvolume. Preprocessing pipelines followed the overall outline of Tinnermann et al. (2022) ^21^.

The first two volumes were removed to eliminate T1 saturation effects. Brain data preprocessing was performed using SPM12 (Wellcome Trust Centre for Neuroimaging, London, UK) and Matlab2022b, spinal data processing was implemented using the Spinal Cord Toolbox ^66^.

### Brain image preprocessing

T1-weighted anatomical images were coregistered to the T1-weighted anatomical MNI152 template from the cat12 toolbox to adjust for the low isocenter positioning near vertebrae C2/C3. EPI images were then coregistered to the T1-weighted anatomical MNI152 template by applying the same parameters, followed by masking using the ArtRepair toolbox to reduce potential ghosting artefacts. Next, all EPI images underwent rigid-body motion correction with six degrees of freedom. The mean EPI image was then coregistered to the anatomical image, segmented, and spatially non-linearly normalized to the Montreal Neurological Institute (MNI) standard space using the cat12 T1-weighted template and Dartel. Following normalization, all functional brain images were warped using individual Dartel flow fields and smoothed with a 6 mm full-width-at-half-maximum (FWHM) isotropic Gaussian kernel.

### Spine image preprocessing

The spinal cord image preprocessing was performed using the Spinal Cord Toolbox. The spinal cord in the T1-weighted anatomical image was initially segmented using a deep learning model. To enhance normalization quality, the segmented image was smoothed, and segmentation was repeated on the smoothed image. Vertebral bodies were then automatically labeled, with manual initialization at the C2/C3 vertebral levels performed if automatic labeling failed (n=14). The T1 image was subsequently normalized to the PAM50 T1 template through a series of steps using the *sct_register_to_template* function: straightening the image using the spinal cord segmentation, aligning vertebrae between the image and template, and applying iterative slice-wise non-linear registration. The PAM50 T2 template was warped into native space using the T1 reverse warp field, as EPI-to-anatomical image registration in native space is more robust when using a T2 reference image compared to a T1 image.

The first two volumes of the EPI images were discarded. The remaining EPI images underwent motion correction using slice-wise registration regularized along the Z direction, and a mean image per task was calculated. Registration to the first task’s mean image (resting state) was performed using affine transformations, followed by computation of an overall mean EPI image. This overall mean was segmented, and the segmentation was visually inspected and manually corrected when necessary. The spinal cord segmentation mask was then applied for registration of the mean EPI image to the T2 image in native space. Finally, the coregistered mean EPI image was normalized to the PAM50 T1 template initialized by the warping and inverse warping fields derived from T1 anatomical-to-template normalization.

Three concatenated warping fields were applied to all remaining EPI images: task mean image (acquisition / extinction) to the first task mean image (resting state), overall mean EPI image to the PAM50 T2 template in native space, and overall mean EPI image to the PAM50 T1 template. All preprocessing steps, including motion correction, coregistration, and normalization, were carefully reviewed through visual inspection to ensure accuracy.

### First level analysis

#### fMRI brain analysisx

Statistical analysis of fMRI data was conducted in SPM12 using a general linear model (GLM) for the brain subvolume. The design matrix for the acquisition task included the following regressors of interest: CS+ non-reinforced, CS-, and CS+ reinforced trials (boxcar trial duration), each with trial-by-trial heart rate (HR) change as a parametric modulator.

Additionally, a regressor for shock delivery (event related) was incorporated. The design matrix for the extinction task included regressors of interest CS+, and CS-trials, each with trial-by-trial heart rate (HR) change as a parametric modulator. Nuisance regressors included six motion parameters, physiological noise regressors based on the RETROICOR method (Glover et al., 2000, https://github.com/can-710lab/RETROICORplus) with respiratory and cardiac harmonics (26 regressors in total), the mean signal within a cerebrospinal fluid (CSF) mask for each volume, and one regressor for volumes exceeding a mean voxel intensity greater than 3 standard deviations from the overall mean voxel intensity of the task data. High-pass filtering was applied with a 1/128 Hz cutoff, and serial correlations were corrected using an AR(1) model.

Event-related responses of regressors of interest were modeled by convolving a 6-second boxcar function with a canonical hemodynamic response function (HRF); the shock regressor was modeled by convolving a stick function with the HRF. Contrasts were defined to identify regions showing differences in activation between CS+ and CS-, as well as HR-modulated activation differences for CS+ and CS-.

#### fMRI spinal cord analysis

For the spinal subvolume, a more temporally precise finite impulse response (FIR) analysis was implemented. The analysis for acquisition and extinction included the same regressors of interest, parametric modulators trial-by-trial HR change, two motion regressors (axial x,y translation) and nuisance regressors as used in the brain analysis. Low-frequency fluctuations from CSF were removed using principal component analysis by modeling components that explained 90% of signal variance as nuisance regressors ^21,48,68^. Regressors of interest (CS+ non-reinforced, CS-) were modeled in six time bins of TR length (3.02 s), covering one TR before stimulus onset to two TRs after stimulus offset (total duration: 18.12 s).

Beta estimates for HR change modulated spinal cord activity were extracted using PAM50 masks for grey matter, dorsal, and ventral regions. The data were then modeled using a Bayesian ANOVA framework in RStudio (R version 4.2.2) **brms**, with condition (CS+ vs. CS-) and time as factors. Priors were specified as normal distributions (μ=0, σ=5), and model fitting was conducted using a skew-normal likelihood with four Markov chains (6000 iterations each, 3000 warmup). Posterior distributions were used to test hypotheses about condition effects over time, and credible intervals were calculated to assess the reliability of effects. Highest Density Intervals (HDIs) were calculated to summarize the posterior parameter distributions, providing a measure of uncertainty. A 95% HDI indicates the range containing 95% of the most credible parameter values. If a 95% HDI excludes 0, the effect can be considered "significant"; if a 90% HDI excludes 0 but the 95% HDI does not, the effect can be considered "marginally significant."

### Second level analyses

Second-level analyses were conducted to examine group-level contrasts between conditioned stimuli (non-reinforced CS+ and CS-) using SPM12. The analyses aimed to evaluate differences in neural responses to CS+ and CS-within the brain subvolume. First-level contrasts calculated for each participant were entered into a one-sample t-test design and analyzed using a factorial design with standard global normalization and masking parameters. One-tailed t-contrasts were defined to test for both CS+ versus CS-activation and deactivation. These analyses were repeated for HR change modulated CS+ and CS-differences. Whole-brain analysis inferences were made at the cluster level (*pFWE <* 0.05), based on a cluster-forming threshold of *p <* 0.005 (uncorrected).

### Connectivity analysis

#### Psychophysiological Interaction (PPI) Analysis

A post hoc exploratory PPI analysis was performed to examine functional connectivity between the spinal cord and brain during acquisition. A general linear model (GLM) was first applied to the spinal cord data, using the same regressors of interest and nuisance regressors as in the FIR analysis. Event-related responses for regressors of interest were modeled by convolving a 6-second boxcar function with a canonical hemodynamic response function (HRF). Time series were then extracted from the grey matter, dorsal, and ventral spinal regions (probabilistic segmentations, threshold 0.5) and adjusted for variance explained by all nuisance regressors. A PPI was created separately for each spinal VOI, based on the contrast of HR-modulated responses between CS+ and CS-trials.

The resulting PPI terms were incorporated into a second-level brain GLM to identify regions functionally coupled to the spinal ROIs. This model included three regressors: the interaction term (PPI), the VOI signal (physiological regressor), and the task contrast (psychological regressor: CS+HR > CS-HR), along with all nuisance regressors. Contrast images were generated for the PPI interaction regressor testing for coupling changes between the heart rate modulated spine (CS+HR > CS-HR) and brain activity.

We pre-defined two ROIs informed by the second-level brain subvolume analysis contrast without HR modulation (peak voxel coordinates of cingulate with x=0, left motor cortex, sphere 10mm) as well as PAG (^16^ sphere 10mm).

Whole-brain analysis inferences were made at the cluster level (*pFWE <* 0.05), based on a cluster-forming threshold of *p <*0.005 (uncorrected). For ROI analyses, a small volume correction (SVC; *p <* 0.05) was performed.

---

**Extended Data Figure 1.**
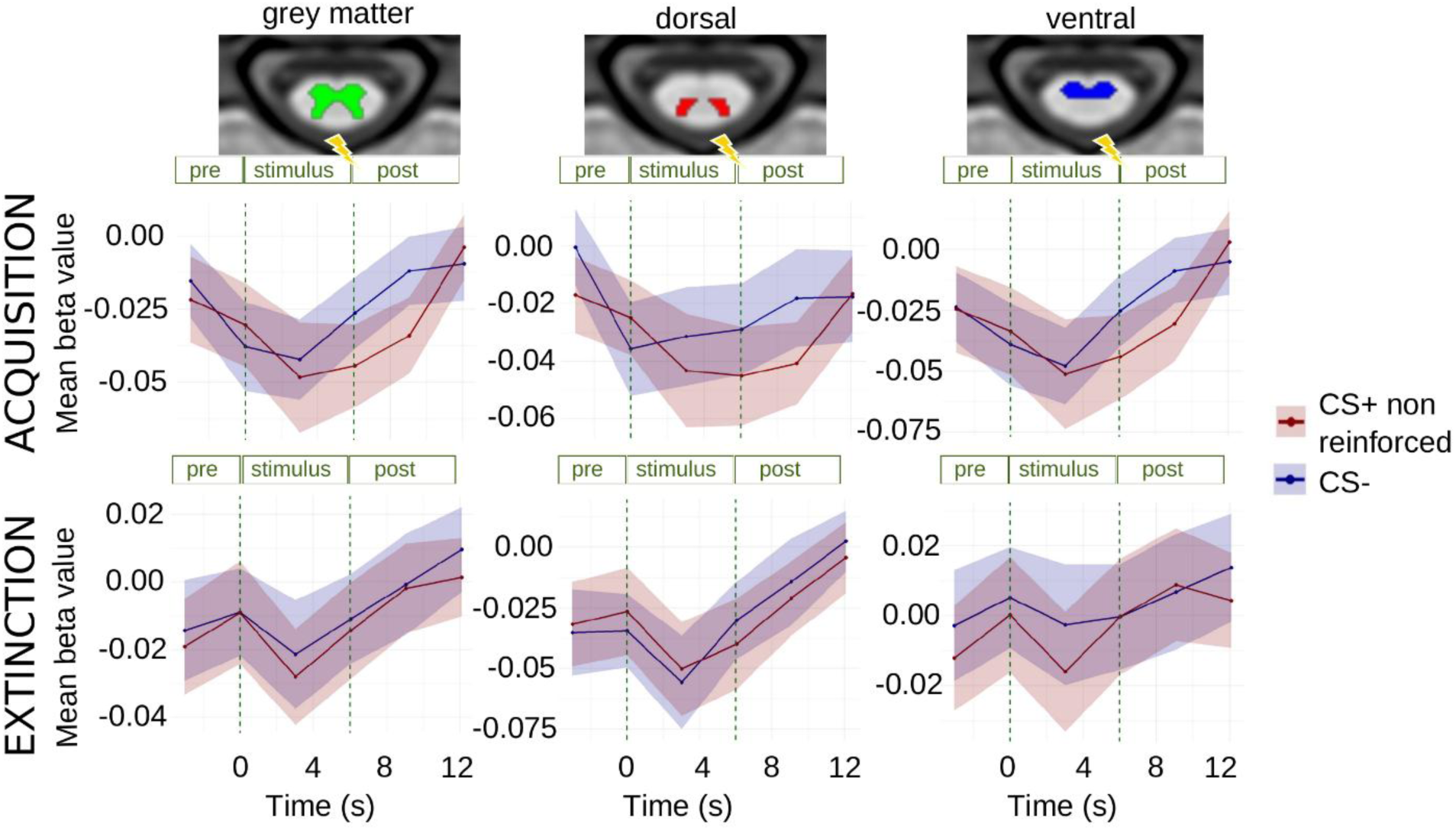
Spinal activity towards CS+ and CS- in a time-resolved FIR analysis. Spinal cord grey matter (left), dorsal (mid) and ventral (right) responses without HR modulation starting one TR before stimulus onset revealed no significant differences in CS+ responses compared to CS- responses in conditioning and in extinction.

**Extended Data Figure 2.**
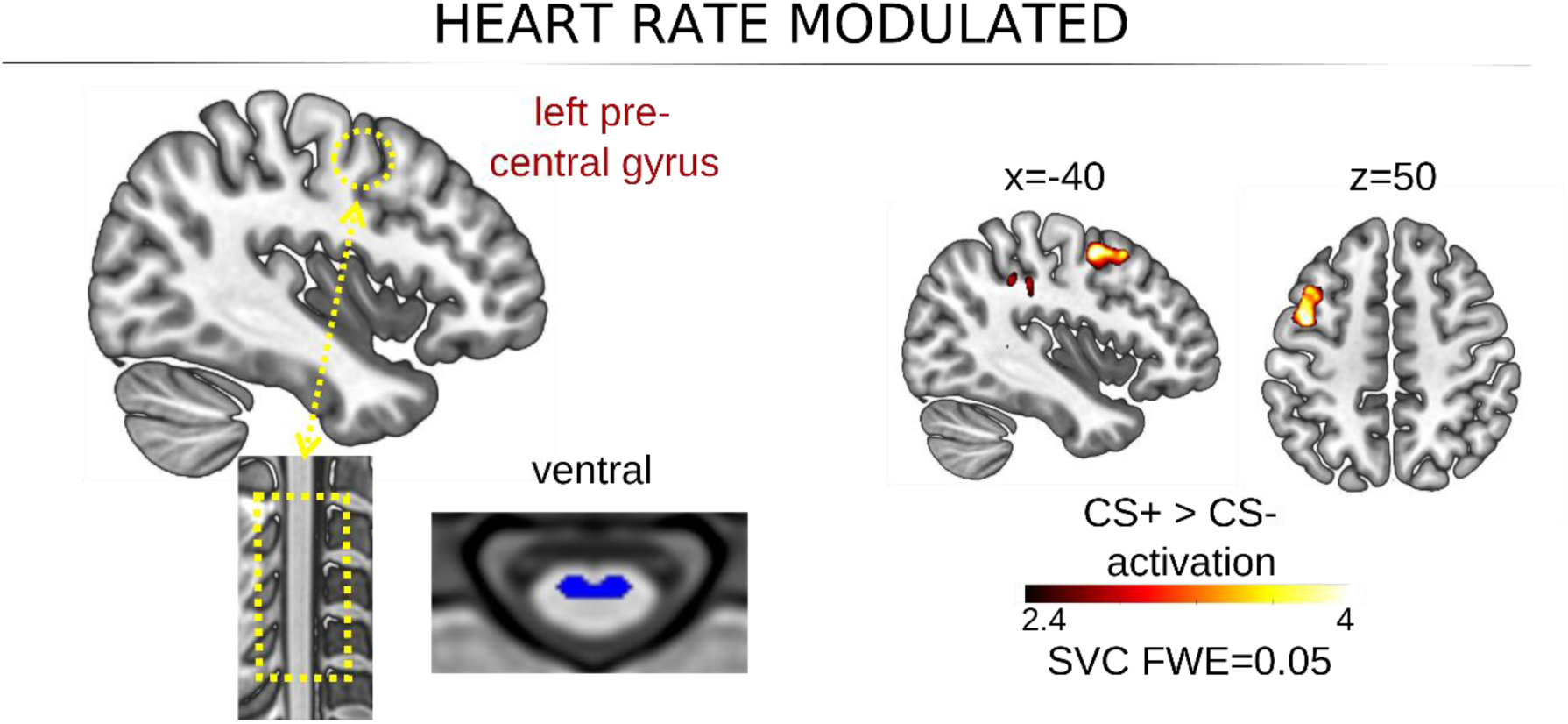
HR-modulated coupling between ventral spine and motor cortex at CS offsets. A PPI based on a GLM using CS offsets during acquisition (timepoints of significant HR- modulated FIR CS+ vs. CS- difference in spinal cord) taking the ventral spinal cord as a seed revealed coupling to the left precentral gyrus post-stimulus offset. SVC = small volume correction applied.

## Notes

### Competing Interest Statement

The authors have declared no competing interest.

